# A rare recombination allows the redefinition of a major avirulence gene in the *Phytophthora sojae* genome

**DOI:** 10.64898/2025.12.17.694715

**Authors:** Yanick Asselin, Caroline Labbé, Vanessa Tremblay, François Belzile, Richard Bélanger

## Abstract

Significant yield losses in soybean production are imputable to Phytophthora root and stem rot (PRR) caused by *Phytophthora sojae*. Soybean resistance to the pathogen depends on the presence of resistance genes (*Rps*) that recognize key effectors *from P. sojae*, which are encoded by avirulence genes (*Avr*). A unique molecular signature associated with these genes allows the prediction of the outcome of infection with great accuracy making the interaction *Rps*-*Avr* central to reduce disease incidence. In this study, we reassessed the identity of the avirulence gene whose protein product is recognized by *Rps6*. Following extensive soil sampling and single-spore isolation of a large population of *P. sojae*, we found two salient isolates carrying a rare recombination between two effectors, *Avr3c* and *Avr4/6*. Using a PCR assay and molecular markers, we showed that only alleles at the *Avr3c* locus were in perfect association with the phenotypes of the isolates. Furthermore, whole-genome resequencing and *de novo* assembly of the two isolates revealed the full extent of this genomic rearrangement. These results bring to light an unsuspected connection between *Avr3c* and *Rps6* and offer a more reliable target for the pathotyping of *P. sojae*, which ultimately leads to a better use of resistant soybean material.

## Main text

*Phytophthora sojae* (Kauf. & Gerd.), the causal agent of Phytophthora root and stem rot (PRR), is one of the ten most important oomycete plant pathogens (Kamoun et al., 2015). It caused about 200 million dollars of yield losses in the United States alone in 2024 (Network, 2025). Management of PRR relies predominantly on the use of genetic control where major Resistance to *Phytophthora sojae* (*Rps*) genes offer reliable protection from germination to harvest. Over 40 *Rps* genes and alleles have been reported (Chen et al., 2025) but only six are commercially exploited, *Rps1a, Rps1b, Rps1c, Rps1k, Rps3a* and *Rps6*. For the optimal control of PRR, the deployment of an *Rps* gene must overcome the *P. sojae* pathotypes present in a field, therefore the specificity of each resistance gene must be precisely known. From the 456 RxLR effector genes revealed by the recent telomere-to-telomere assembly of the *P. sojae* genome (Zhang et al., 2024) only a few selected members are recognized by the product of an *Rps* gene; these include *Avr1a* (Qutob et al., 2009), *Avr1b* (Shan et al., 2004), *Avr1c* (Na et al., 2014), *Avr1d* (Na et al., 2013), *Avr1k* (Song et al., 2013), *Avr3a/5/8* (Arsenault-Labrecque et al., 2022; Dong, Yu, et al., 2011; Qutob et al., 2009), *Avr3b*/*11* (Asselin et al., 2025; Dong, Yin, et al., 2011), *Avr3c* (Dong et al., 2009) and *Avr4/6* (Dou et al., 2010). Distinct SNP haplotypes have been associated with alleles of each *Avr* gene to predict the response to infection of resistant soybean lines (Arsenault-Labrecque et al., 2018). The near-perfect association between *Rps* genes and their corresponding effector makes this pathosystem a salient example of the gene-for-gene concept based on effector-triggered immunity (ETI).

In spite of their initial distinction, *Rps4* and *Rps6* have been shown to recognize *Avr4* and *Avr6*, two genes originally described as tightly linked but later renamed *Avr4/6* because of the complete cosegregation of the traits (Gijzen et al., 1996; Whisson et al., 2004). Crosses between *P. sojae* isolates, discriminant for their virulence against *Rps4* and *Rps6*, allowed the mapping of the avirulence gene on chromosome 5 (Chr5) in a recombination-rich region close to another major avirulence gene, *Avr3c* (Dong et al., 2009; Dou et al., 2010). A transient expression assay showed that both *Avr4/6* and *Avr3c* induced cell-death in resistant soybean lines. However, in this study, we gathered evidence of a rare recombination between the two closely linked genes coding for these effectors. Molecular markers were employed in a PCR assay to show that only alleles at the *Avr3c* locus were associated with virulence towards *Rps4* and *Rps6*. Furthermore, long-read whole-genome resequencing and *de novo* assembly showcased the rearrangement that redefines the identity of a major avirulence gene.

During a three-year campaign in five Canadian provinces, soil samples were harvested in soybean farms with a history of PRR presence. Single-spore isolation was performed on the soils according to the baiting method described by Tremblay et al. (2021). A collection of 236 isolates was retrieved and fully pathotyped using a differential of 13 resistant soybean NILs and one susceptible control in a hydroponic assay (Lebreton et al., 2018). All isolates were virulent on the susceptible control and showed a wide range of pathotypes (Table S1.) From the phenotyping experiment, 31 of the 236 isolates were capable of infecting both *Rps4* and *Rps6* resistant soybean lines, with no isolate virulent to only one line, representing an occurrence of 13.1%. All remaining isolates (205) were avirulent to these same lines but virulent against other resistant lines. Pure cultures were grown on V8 agar media covered by cellulose to facilitate mycelium harvest. Then, samples were homogenized and DNA was extracted using a modified CTAB method. Using the sequence of *Avr3c* published by Dong et al. (2009), PCR primers were designed to discriminate virulent and avirulent alleles of the gene. For *Avr4/6*, no variation is present in the coding regions of the virulent and avirulent alleles, but a 15-bp deletion upstream was found to be associated with virulence according to Dussault-Benoit et al. (2020). This polymorphism was therefore employed as a marker to distinguish these alleles (Table S2.). All 236 isolates were then genotyped for their alleles of *Avr3c* and *Avr4*/6 in a PCR assay. Amplification yielded results in all samples meaning no deletions of these genes were found in the collection. For 234 *P. sojae* isolates, the genotypes of both genes were found to coincide as either virulent or avirulent. For all those isolates, their phenotypes corresponded to their genotypes. However, discrepancies were observed in two isolates. Haplotypes of ULPS-556 for *Avr3c* and *Avr4/6* were respectively avirulent and virulent but the phenotype of the strain was avirulent against *Rps4* and *Rps6*. For isolate ULPS-778, there was an amplification of the virulent allele of *Avr3c* and the avirulent allele of *Avr4/6*. In this case, the phenotype of the strain indicated virulence against *Rps4* and *Rps6*. For ULPS-556 and ULPS-778, a recombination between the two genes occurred and only the genotypes of *Avr3c* corresponded with the phenotypes of the isolates. To validate these results, a modified version of the hypocotyl-wound inoculation from Dorrance et al. (2008) was also performed on Williams, L85-2352 (*Rps4*), L89-1581 (*Rps6*), Harosoy, Haro4272 (*Rps4*) and Haro6272 (*Rps6*)(Figure 1.). Results were consistent with those of the hydroponic assays, thus confirming the sole association of *Avr3c* with virulence towards *Rps4* and *Rps6*.

**Figure 1.**
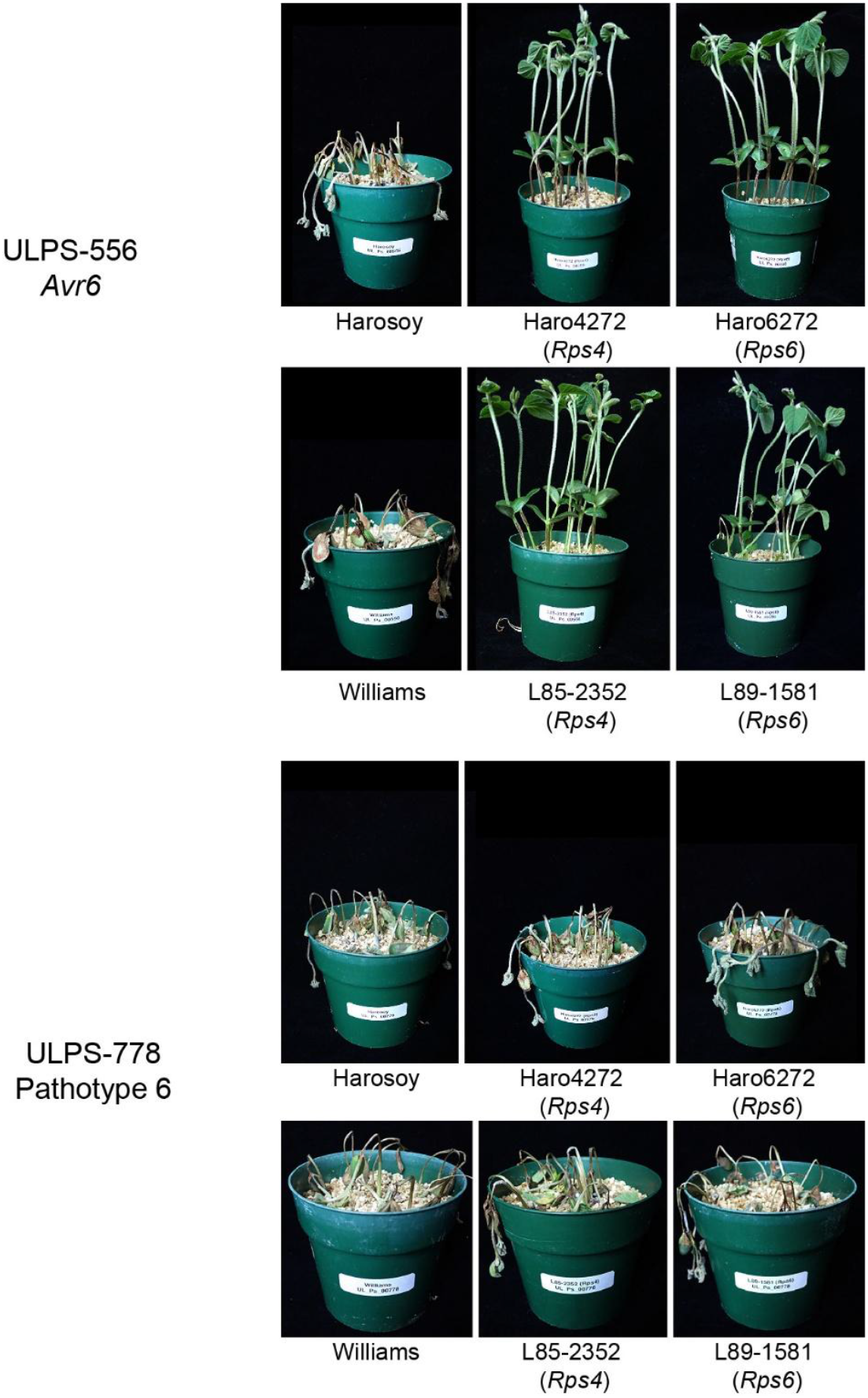
Phenotypic responses of *Rps4* and *Rps6* soybean lines inoculated with *Phytophthora sojae* isolates carrying the virulent or avirulent form of *Avr3c*. Phenotypes of L85-2352 (*Rps4*), Haro4272 (*Rps4*), Haro6272 (*Rps6*), L89-1581 (*Rps6*) and susceptible controls (Williams and Harosoy) using the hypocotyl-wound inoculation. Isolate ULPS-556 carries the avirulent allele of *Avr3c* and isolate ULPS-778 carries the virulent one.

To investigate this rare recombination between effectors *Avr3c* and *Avr4/6* in isolates ULPS-556 and ULPS-778, sequencing on the MinIon from Oxford Nanopore Technologies (Oxford, UK) and Illumina NovaSeq (San Diego, U.S.) platforms were performed. For ULPS-556, coverages of 68X and 158X were obtained from long- and short-read sequencings, respectively, whereas the sequencing of ULPS-778 generated 46X and 150X of data. For each isolate, the local *de novo* assembly of the *Avr3c* and *Avr4/6* region on Chr5 produced gap-free contigs of 175 kb and 180 kb including 50 kb up- and downstream of the genes. For comparative purposes, two other isolates without the recombination between the two effectors were also sequenced, one virulent to *Rps4* and *Rps6* (ULPS-728) and one avirulent (ULPS-733). Assemblies, from long and short reads, of 175 kb and 213 kb were respectively produced as virulent and avirulent references and annotation was produced using the latest *P. sojae* genome assembly (Zhang et al., 2024). Global synteny, between physical position 343,700 and 518,850 of Chr5, was evaluated among the four isolates using MCScanX (Wang et al., 2012). As shown in Figure 2, the recombinant isolate ULPS-556 shows syntenies with both the avirulent and virulent references. As a matter of fact, the *Avr4/6* gene and its haplotype display overall greater sequence identity with the virulent reference except for the avirulence gene itself because no variants are found in its coding sequence. In comparison, the *Avr3c* region of ULPS-556 is nearly identical with the avirulent reference as almost no SNPs are observed. The mirror image is observed for isolate ULPS-778, where the *Avr4/6* region matches the avirulent reference and *Avr3c* the virulent one. Once again, no variants were found within the coding sequence of *Avr4/6*. The synteny analysis showed a higher structural conservation around *Avr4/6*; except for SNPs and a few small indels, no major structural rearrangements were found. However, a different scenario is detected around *Avr3c*. We observed a segmental duplication of a block of 33.7 kb, as reported by Dong et al. (2009), in the isolates avirulent to *Rps4* and *Rps6*. Three copies were found in our avirulent reference (ULPS-733) but only two in ULPS-556. The virulent *P. sojae* isolates also carried a repetitive segment, but one of 29.2 kb, with a haplotype distinct from the avirulent one. Altogether, greater variation is observed throughout the *Avr3c* region whereas *Avr4/6* is more conserved.

**Figure 2.**
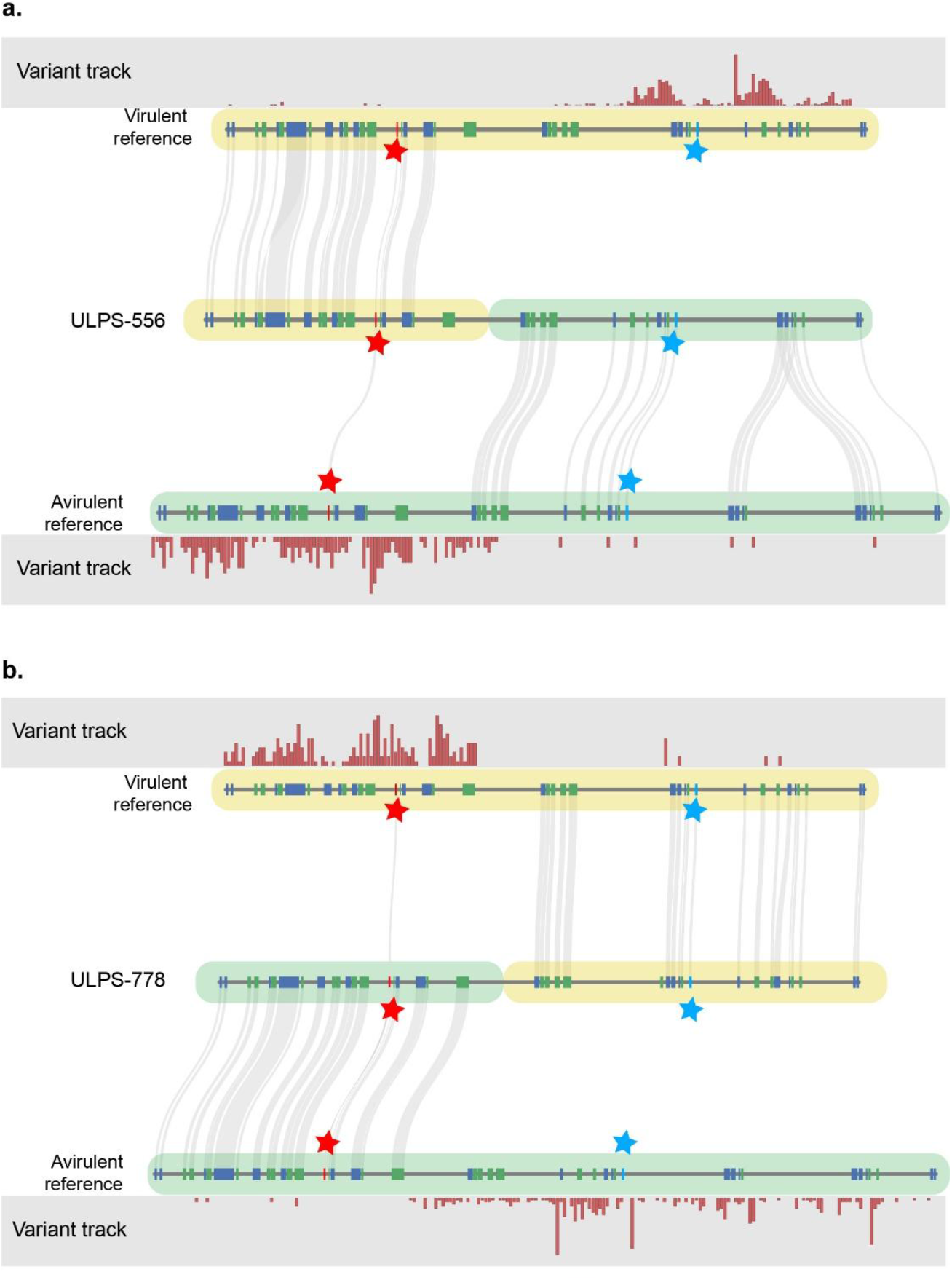
Recombination between the *Avr4/6* and *Avr3c* avirulence gene in two *Phytophthora sojae* field isolates. Synteny on Chr5 from positions 343,700 to 518,850 and variant coverage among the avirulent isolate ULPS-556 (a) and the virulent isolate ULPS-778 (b) compared to a virulent (ULPS-728) and an avirulent (ULPS-733) references. The green and yellow shaded areas represent the avirulent and virulent haplotype. Green and blue arrow boxes represent the orientation of the predicted coding sequences of all genes and the grey shaded areas represent conserved syntenic blocks. The red and blue stars mark the location of *Avr4/6* and *Avr3c* respectively.

Plant pathogens exhibit tremendous intraspecific genetic variation, especially for genes under strong selection pressure to overcome host resistance. Among those, effectors and more specifically avirulence genes are offering some of the most diverse regions in their genomes. In this study we found a rare recombination in the *Phytophthora sojae* genome that allowed to reassess the specificity of two major resistance genes, *Rps4* and *Rps6*. While one avirulence gene, *Avr4/6*, has been previously associated with this source of resistance, we present evidence that the recognition process is rather linked with another one, *Avr3c*. Interestingly, based on a large natural population of *P. sojae* isolates, we showed that alleles of *Avr3c* and *Avr4/6* were in perfect linkage in 234 out of 236 isolates. Original mapping by Whisson et al. (2004) positioned *Avr4/6* in a 4.3 cM region of Chr5. Assuming a 35.3 kb/cM recombination rate (Whisson et al., 2004), the 4.3 cM can possibly cover up to 151.8 kb, thereby including the *Avr3c* locus. We therefore suggest that Whisson et al. (2004) and Dong et al. (2009) both correctly associated the same locus with virulence towards resistant soybean lines

*Phytophthora sojae* can evade recognition from its host by exploiting diverse strategies such as the mutation of key residues in the protein produced by its avirulence genes as in the case of *Avr1c, Avr3b* and *Avr3c* (Huang et al., 2019; Kong et al., 2015; Yang et al., 2019). Complete deletion of the genes *Avr1a, Avr1b, Avr1c* and *Avr1d* has also been described as another route for promoting virulence of the pathogen (Cui et al., 2012; Na et al., 2014; Na et al., 2013; Qutob et al., 2009). Finally, gene silencing by siRNAs was described for *Avr3a* but was also proposed for *Avr1a* and *Avr1c* (Qutob et al., 2013; Qutob et al., 2009; Yang et al., 2019). Considering all putative evasion strategies, *Avr4/6* is an oddity among avirulence genes. According to Dou et al. (2010) no sequence polymorphisms are present in its coding sequence and no transcriptional changes between virulent and avirulent strains are measured. This raises the question as to how can this protein avoid being recognized? Based on our results, it appears that *Avr4/6* does not act as a *bona fide* avirulence gene. While it was shown to induce cell death in *Rps4* and *Rps6* soybean lines, it is unclear whether the recognition was specific to this source of resistance or if the protein also induced immune responses in other resistant genotypes. Indeed, in *Phytophthora* species, some conserved effectors can trigger cell death without necessarily generating a resistance response in the plant because the virulence mechanisms of a pathogen are an equilibrium involving hundreds of secreted proteins so that the effect of one is not always a good proxy for the whole plant response (Oh et al., 2024).

In this work, we provide new insights into the *P. sojae* virulence mechanisms and show that *Avr3c*, the corresponding effector of *Rps3c*, is also recognized by *Rps4* and *Rps6*. This result is coherent with the recent report that all three genes are in fact redundant (Asselin et al., 2025) and suggests that *Avr3c* stands as the correct and sole avirulence gene for all three *Rps* genes and should be renamed to better align with *Rps6*, the only one commercially deployed. Knowledge of the specificity of a resistance gene is of paramount importance for the durable control of Phytophthora root and stem rot. Recent field surveys around the globe showed that *Rps6* is still largely effective (McCoy et al., 2023) and, to maintain its efficacy, an integrated approach involving producers and agronomists, must result in an effort to characterize the *P. sojae* populations so that resistant cultivars are only deployed where they are effective.

## Acknowledgments

This research was supported by Genome Canada and the Canada Research Chair in plant protection of Richard R. Bélanger.

## CRediT author contributions

Y.A.: Conceptualization; data curation; formal analysis; methodology; validation; writing original draft. C.L.: Investigation, Methodology; project administration; supervision. V.T.: Investigation. R.R.B.: Conceptualization; funding acquisition; project administration; supervision; writing review and editing.

